# Comparative genome-scale constraint-based metabolic modeling reveals key lifestyle features of plant-associated *Pseudomonas* spp

**DOI:** 10.1101/2022.07.26.501552

**Authors:** Wasin Poncheewin, Anne D. van Diepeningen, Theo AJ van der Lee, Peter J. Schaap, Vitor A. P. Martins dos⍰Santos, Maria Suarez-Diez

## Abstract

Plant Growth Promoting Rhizobacteria (PGPR) dwell in the rhizosphere, the area surrounding the root of plants, and enhance growth of the host through different mechanisms: they can protect plants against pathogens, assist in nutrient gathering, and in increasing stress tolerance. Hence, developing strategies to enhance their performance is important to increase crop productivity. Specific solutions are necessary to enhance the performance of the beneficials while simultaneously avoiding nurturing of pathogens. This requires insights into the mechanisms underlying these microbials interactions. *Pseudomonas* is one of the most studied genera and contains both beneficials and pathogenic species. Hence, we used comparative genome-scale constraint-based metabolic modeling to reveal key features of both classes of Pseudomonads and which can provide leads for the possible interventions regarding these solutions. Models of 75 plant-growth promoting rhizosphere and 33 epiphytic pathogenic *Pseudomonas* strains were automatically reconstructed and validated using phenotype microarray (Biolog) data. The models were used for compositional analysis and 12 representative strains, 6 of each group, were further selected for extensive simulation. The analyses reveal differences in the potential for metabolite uptake and transport between these two distinct classes that suggest their nutrient preferences and their differences in, among other, D-ornithine acquisition mechanisms. The models enable simulation of metabolic state of root exudates. Simulations highlighted and summarized the differences in pathway utilization and intracellular states between two groups. The insights obtained will be very valuable to broader such studies of rhizobiome and to possibly develop strategies to improve crop productivity by supporting the beneficial microbiome while reducing pathogen activities.

## Introduction

The rhizosphere, the interface between the plant root and the soil, is influenced by the chemicals released from the plant root system and can be inhabited by a population of plant beneficial microorganisms and sometimes pathogens attracted by such plant exudates^1,2^. The attracted beneficial organisms benefit their host by enhancing nutrient acquisition and tolerance to biotic and abiotic stresses^3–5^. Bacterial rhizosphere community members are often represented by a diverse set of taxa often with Pseudomonadaceae as one of the predominant groups^6–8^.

The most studied genus within the Pseudomonadaceae is the name-giving genus *Pseudomonas*. Members of this genus vary in lifestyle, organic compound utilization and habitation of ecosystems and the genus contains both plant beneficial and plant pathogenic species^7–9^. Most of the plant beneficial *Pseudomonas* strains identified belong to the *P. fluorescens* species group, while most of the identified plant pathogens are *P. syringae* strains^10,11^. However, there are exceptions such as the plant growth promoting *P. syringae* pv. *syringae* strain 260-02^12,13^. This suggests that the functional significance or biochemical role of a given strain in a defined environment such as the rhizosphere can potentially be prioritized over taxonomy^14–16^. Moreover, the dynamic environment that accommodated these microbes compels them to adapt to changes for their own and host survivability^17,18^. For these reasons, the investigation of the metabolic differences of two distinct classes, beneficial and pathogen, can reveal the common unique characteristics per group.

Genome data is available from many environmental isolates and genome-scale metabolic network reconstructions (GEMs), coupled with constraint-based analysis methods and tools, such as Flux Balance Analysis (FBA)^19,20^, allow the comparison of their metabolism and transport at a systems-level. Such comparative studies are vital to understand the principles and mechanisms involved in defining the specific traits contributing to a plant beneficial or pathogenic phenotype.

In this study we utilized the CarveMe automatic GEMs reconstruction tools^21,22^ to compare GEMs from 75 known Plant-Growth Promoting Rhizobacteria (PGPR) with 33 Epiphytic Pathogenic *Pseudomonas* (EPP) strains originating from various *Pseudomonas* spp. using new and available genome sequences and a standardized de novo annotation pipeline as input^23^. This allowed us to elucidate systems-level metabolic differences between the two classes and by simulating different environmental conditions, in a time-series manner, medium specific reactions were revealed^24^. The results show that GEMs can identify different nutrient preferences through the annotated transports and pinpoint differences in pathway wiring towards optimal growth. This crucial knowledge can be implemented further to enhance crop productivity by precisely assisting the beneficial microbiome while reducing pathogen activities.

## Materials and Methods

### Genome retrieval and annotation

Genomes of seven beneficial *Pseudomonas* strains: *P. putida* P9 (accession ERS6670306), *P. corrugata* IDV1 (accession ERS6652532), *P. fluorescens* R1 (accession ERS6670181), *P. protegens* Pf-5 (accession ERS6652530), *P. chlororaphis* Phz24 (accession ERS6670416), *P. jessenii* RU47 (accession ERS6670307) and *P. fluorescens* WCS374 (accession ERS6652531) have recently been re-sequenced^25^. In addition, 101 publicly available “complete” *Pseudomonas* genomes were downloaded from the *Pseudomonas* Genome DB version 20.2 (https://www.pseudomonas.com)^26^. The downloaded data were categorized according to the literature into two classes: Plant-Growth Promoting Rhizobacteria (PGPR) (68 strains) and Epiphytic Pathogenic *Pseudomonas* (EPP) (33 strains). These sequences were annotated with protein domains and synteny-non directional Genome Properties (SND-GPs). The annotated data along with their literature references of the complete and classified genome were obtained from Poncheewin et al.^25^.

### Model construction

CarveMe v.1.5.1 was used to automatically construct gap-filled genome scale metabolic models (GEMs) from the annotated protein domains using aerobic M9 minimal medium with the universal template of the gram-negative bacteria in BiGG models^21,27^ as growth conditions. The availability of metabolites in M9 was simulated by setting the lower-bound of the corresponding exchange reactions, which transfer metabolites in and outside of the organism, to -10 mmol/gDW/h along with the oxygen exchange reaction to simulate the aerobic condition. In the models, the exchange reactions corresponding to the M9 medium components are termed EX_glc__D_e, EX_o2_e, EX_ca2_e, EX_cl_e, EX_cobalt2_e, EX_cu2_e, EX_fe2_e, EX_fe3_e, EX_h2o_e, EX_h_e, EX_k_e, EX_mg2_e, EX_mn2_e, EX_mobd_e, EX_nh4_e, EX_pi_e, EX_so4_e and EX_zn2_e and are used to simulate availability of the corresponding components.

### Model composition analysis

An enrichment analysis was performed on the model’ s reactions. Hypergeometric tests with Bonferroni correction were used on each class to uncover over-representative reactions (p-value < 0.05) using dhyper and p.adjust functions in R^28^. The enriched reaction’s descriptions were used to create a document to illustrate a word cloud using “tm” v.0.7-8, “RColorBrewer” v.1.1-2, “wordcloud” v.2.6 and “wordcloud2” v.0.2.1 packages^29–32^. BLASTP within DIAMOND v.0.9.14.115 was used to obtain the similarity score of the protein sequences related to D-ornithine activity: D-ornithine transport via ABC system periplasm (DORNabcpp) and ornithine racemase (ORNR)^33^. A total of 3 databases were created (1) full set of protein sequences from all the strains, (2) genes annotated to DORNabcpp reaction were removed and (3) genes annotated to ORNR reaction were removed.

### Strain selection

Hierarchical clustering was performed on the statuses of the GPs of each class using Euclidean distance with complete linkage. The functional based dendrograms were pruned using Treemmer v.0.3 for 100 iterations to select 12 representative strains, 6 of each class^34^. Strains with the most frequent occurrences while maintaining the distances of the tree were selected.

### Model simulation

GEMs were used to simulate fluxes and consumption capabilities through Flux Balance Analysis (FBA) using COBRApy version 0.22.0^19,20^. FBA computes reaction fluxes that optimize the flux through a selected objective reaction, which is often selected to represent either growth or production or consumption of a chemical compound of interest. Optimal growth rates were estimated using FBA by setting the biomass synthesis reaction as the objective for maximization. For comparison with Biolog data, the models were used to simulate metabolite consumption profiles. To do so, a total of 55 carbon sources were used to substitute EX_glc_e (glucose) from the initial M9, one at a time The consumption of the tested carbon was limited to 10 mmol/gDW/h by setting the lower bound of the corresponding exchange reactions to -10. The maximum possible consumption of the carbon sources was estimated using FBA by setting the corresponding exchange reactions as minimization objective (**Supplementary file S1**), as negative values indicate consumption of the metabolite. The profile was compared to the Biolog data while the threshold for the ability to oxidize compounds in the Biolog set to 0.1.

Three media were defined to represent different growth stages of the tomato seedling. The M9 media was adjusted by adding additional organic acids and sugars as follow (Day2) M9 with the addition of oxalate (15 mmol/gDW/h) and xylose (11 mmol/gDW/h), (Day4) M9 with the addition of citrate (5 mmol/gDW/h) and fructose (9.17 mmol/gDW/h) and (Day14) M9 with the addition of citrate (5 mmol/gDW/h), xylose (5 mmol/gDW/h) and maltose (2.5 mmol/gDW/h)^24^.

For each medium and model, we performed single reaction deletions to assess their essentiality. The reactions were essential if the growth predicted after deletion was less than 10% of the optimal growth. Reactions were mapped to KEGG PATHWAY for pathway identification using their corresponding EC number yielding in the fraction of completeness per pathways^35^.

We explored growth feasibility and flexibility using 10,000 iterations of flux sampling under parsimoniousFBA (pFBA) constraint towards the optimal growth (**Supplementary file S1**)^36,37^. Flux sampling was performed using optGpSampler as implemented in the COBRApy sample function with “optgp” option ^37^.

### Pathway analysis

Statistical methods were used to identify significant differences in pathways and fluxes between the two classes PGPR and EPP (p-value < 0.05). T-tests were applied to the fraction of completeness of essential pathways between two groups using the t.test function in R^28^. For sampled fluxes, we perform a pairwise comparison between each member of the different groups, resulting in a total of 36 comparisons. Kolmogorov-Smirnov tests (KS test) were applied on the sampled fluxes through the ks.test function in R^38^. In addition to the p-value, the fluxes were considered significant if the distance (D) was more than 0.5 and the mean more than 0.01.

### Metabolic characterization

Biolog phenotyping microarrays were used as suggested by the manufacturer (Biolog, Hayward (CA), USA). Microplates PM1, PM2A, and PM3B were used containing 190 carbon sources and PM10 to test for pH and carbon sources **(Supplementary file S2)**.

## Results

### Model construction and validation

Seven de novo (re)-sequenced and annotated strains: *P. putida* P9, *P. corrugata* IDV1, *P. fluorescens* R1 and WCS374, *P. protegens* Pf-5, *P. chlororaphis* Phz24, and *P. jessenii* RU47 were selected for automatic GEM construction based on M9 minimal medium. The carbon assimilation profile of the models was then simulated and compare with the Biolog data using a carbon substituted M9 minimal medium **(Figure 2)**. Glucose in the M9 medium was substituted with each of the 55 tested carbon sources, one at a time. For each substitution, the carbon source was set as the model’s objective and was minimized to create a carbon assimilation profile per strain which resembles the Biolog data. The comparison allowed us to verify accuracy of the carbon consumption profiles predicted by the models. As the result depicted, the comparison yields 69.35% accuracy with 72.98% and 88.35% for recall and precision respectively.

**Figure 1:**
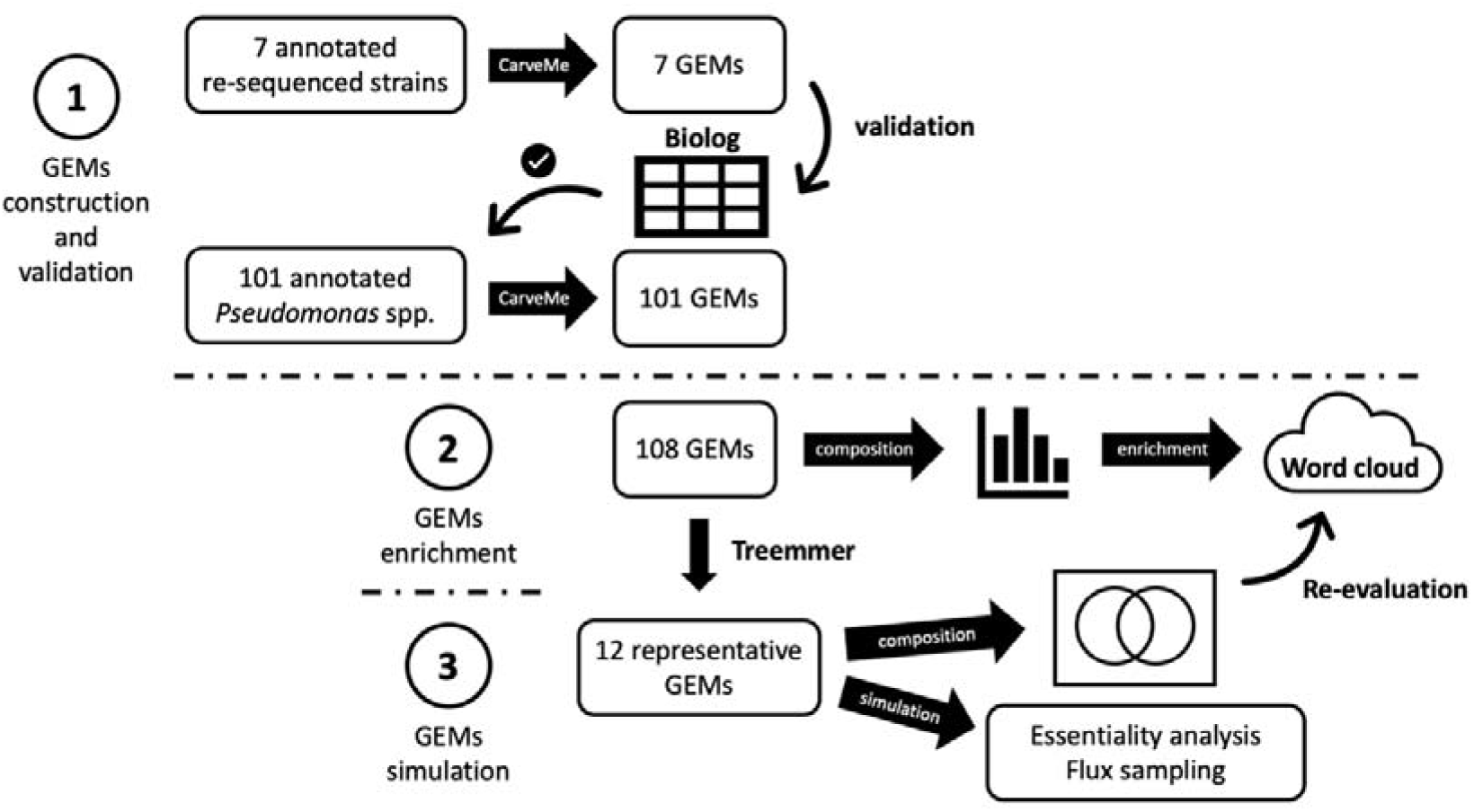
Three step analysis workflow for GEM construction, enrichment, and simulation. Step (1): GEMs representing *P. putida* P9, *P. corrugata* IDV1, *P. fluorescens* R1 and WCS37, *P. protegens* Pf-5, *P. chlororaphis* Phz24, *P. jessenii* RU47 were automatically constructed with CarveMe and validated against the Biolog phenotype data. The validation showed the approach to be suitable and GEMs were automatically built for the 101 remaining strains. Step (2): Reactions from the total set of GEMs were evaluated by enrichment analysis and results were represented with a word cloud. Step (3) Treemmer was used to select 12 representative strains for further in-depth analysis. Corresponding models were explored using enrichment analysis and used for extensive model simulations to identify essential reactions and differences in intracellular fluxes.

**Figure 2:**
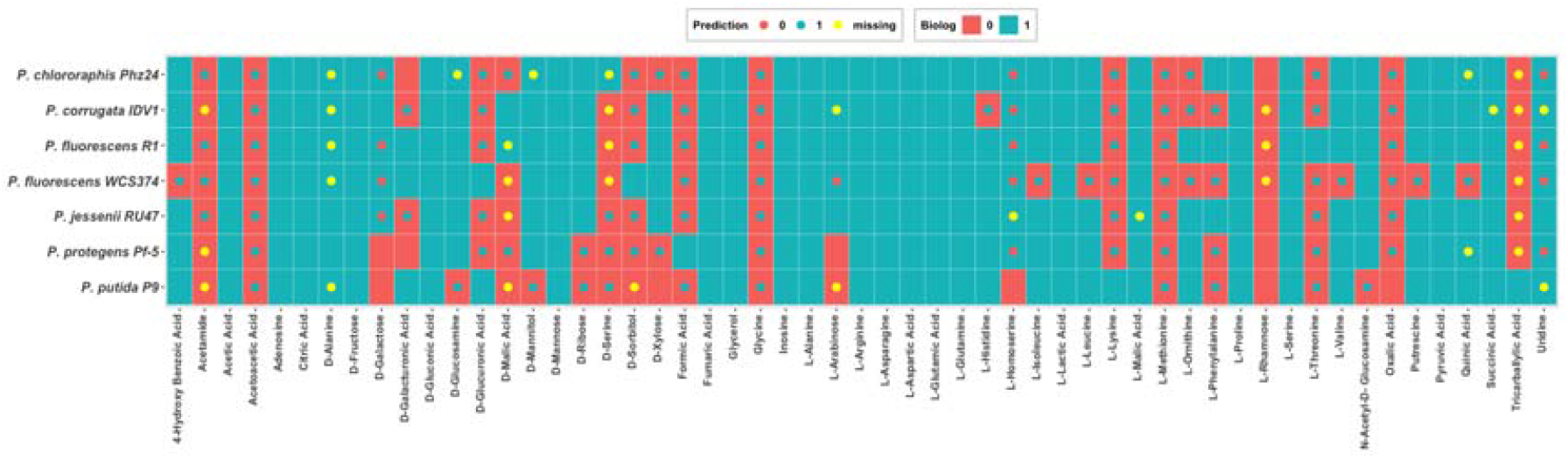
Carbon assimilation profile of the 7 sequenced strains compared to the Biolog data. The square represents the Biolog data while the dot represents the prediction. Blue color (1) represents the ability to oxidize the carbon sources where red (0) is the opposite. The yellow dot are carbon sources that do not exist in the model which were calculated as the inability to oxidize the carbon sources.

Once the performance of the automatic approach was validated, 101 additional GEMs were constructed. The overview of the model composition is summarized in **Figure 3(a) and (b) (Supplementary file S3)**. Figure 3(a) shows the number of genes, metabolites, and reactions. Metabolites are separated by their cellular locations: in the periplasm, extracellular, or in the cytosol. Reactions are categorized into orphan reactions, exchange reactions and reactions with referenced genes (GPRs). In brief, all models are composed of approximately 2,000 genes, 1,700 metabolites and 2,700 reactions. **Figure 3(b)** illustrates a histogram regarding the number of occurrences of reactions excluding the exchange reactions and of cytosol metabolites across all models. Approximately 35% of both reactions and metabolites are shared between all models whereas approximately 5% of both contents are unique to one model.

**Figure 3:**
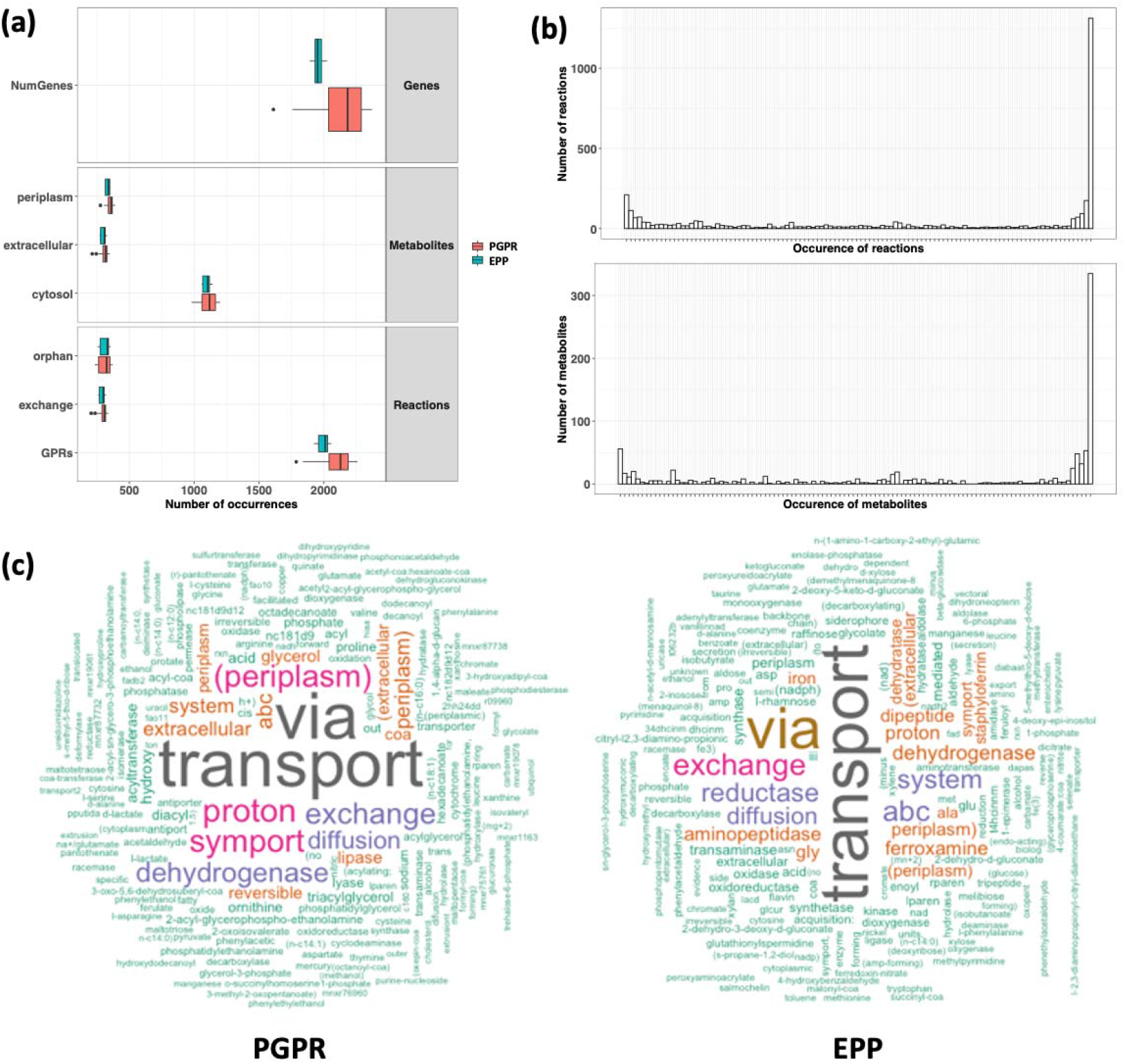
Overview of the model composition. (a) Number of genes, metabolites and reactions separated by the classes. Metabolites are separated by their cellular locations: periplasm extracellular and cytosol. Reactions are categorized into orphan reactions, exchange reactions and reactions with referenced genes (GPRs). (b) Histograms show the number of occurrences of reactions, excluding exchange reactions and cytosol metabolites across all models. (c) Word cloud of the enriched reactions’ descriptions of two classes. The size of the word reflects the number of occurrences of each word.

With more models involved, we assessed differences between classes in their reaction content. As a result, 314 and 197 reactions were found to be enriched in the PGPR and EPP groups, respectively **(Supplementary file S3)**. To summarize the differences, the enriched reactions’ descriptions were visualized using a word cloud **(Figure 3(c)**). Notably with all enriched reactions, the most prominent differences observed are reactions related to transport of metabolites. The PGPRs’ transports were mostly related to amino acid metabolisms, such as alanine, valine, and phenylalanine whereas the EPPs’ transports were annotated with carbon sources, such as galactose, xylose, and sucrose and iron-related metabolites, such as siderophore and staphyloferrin.

### Selection of representative strains

For further in-depth analysis such as flux sampling, which is computationally intensive, we selected representative strains of each class. All sequences were annotated with GPs, and these were used to construct hierarchical trees. The dendrograms were repeatedly pruned down to 6 branches per class and the strains present at the end of the pruning were recorded. After 100 iterations, strains with the most frequent occurrences in the pruned tree while maximizing the distribution of the tree were selected as the representative strains **(Figure 4)**. The six PGPR representatives are *Pseudomonas* sp. UW4 194, *P. chlororaphis* Phz24, *P. fluorescens* WCS374, *P. jessenii* RU47, P. rhizosphaerae DSM 16299 3023 and *P. stutzeri* A1501 123. The six EPP representatives are *P. syringae* ATCC 10859 3811, *P. viridiflava* CFBP1590 isolate E12-5 7308, *P. syringae* pv tomato B13-200 7111, *P. savastanoi* 1448A 114, *P. cerasi* isolate Sour cherry (Prunus cerasus) symptoma 4022 and *P. cichorii* JBC1 2922.

**Figure 4:**
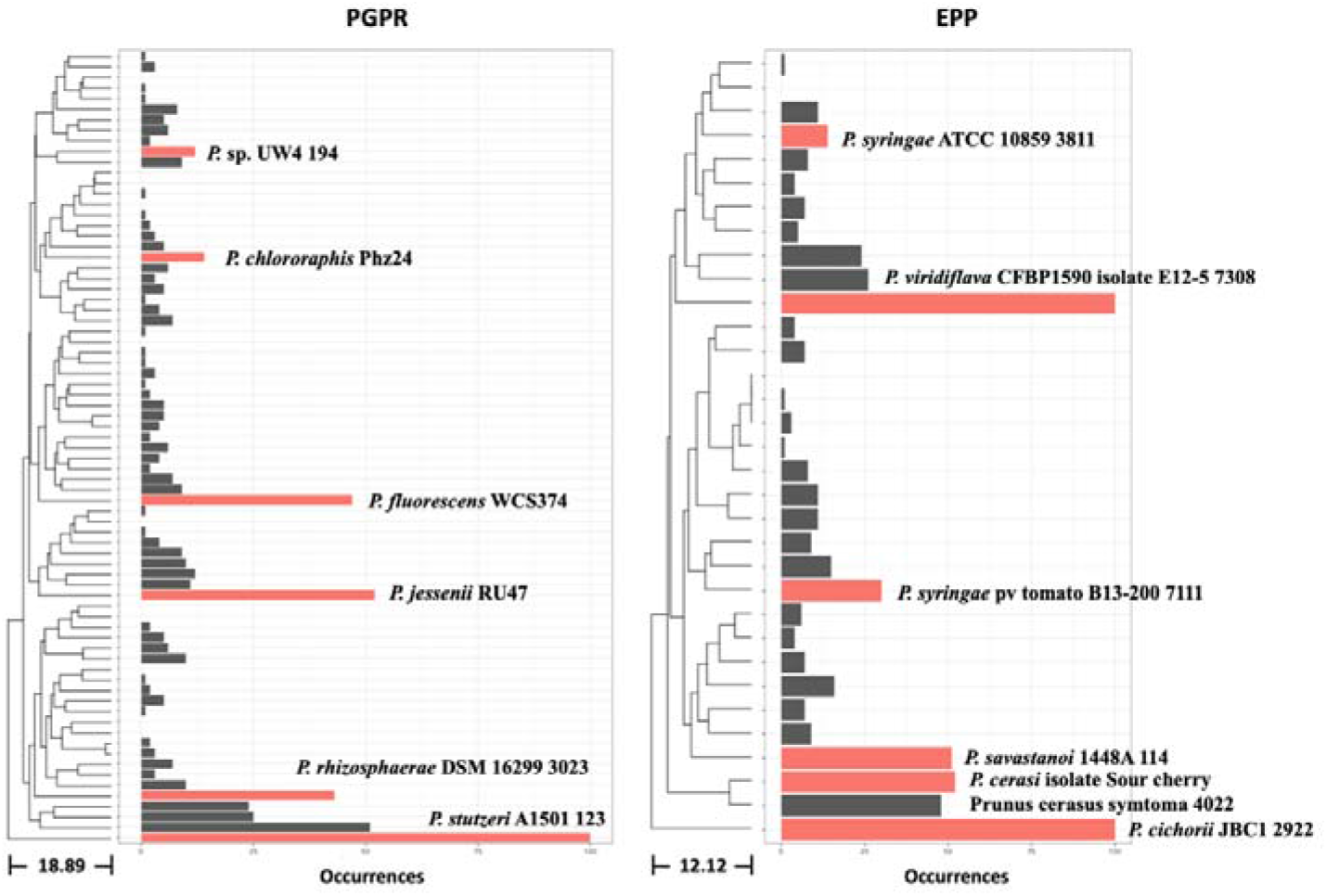
Selection of representative strains. The occurrences of each strain were stored after each prune. The most frequent occurrence strains while maintaining the distribution of the dendrogram were selected as the representative strains.

### Comparative analyses of the metabolic reconstructions

The reactions within the GEMs of the selected strains were compared. Sets of reactions were combined for all models of the same class. We found 3033 reactions were shared between the two classes, whereas 529 and 153 reactions were unique to PGPR and EPP groups, respectively. We further investigated the reactions that were shared between all models within the same group. A total of 4 and 7 reactions were found to be specifically shared within the PGPR and the EPP respectively (**Table 1 and Supplementary file S4**). We also re-evaluate the group-specific reactions with the whole set of constructed models to assess their representativeness on their occurrences in each group along with their adjusted p-value from the enrichment analysis previously performed.

**Table 1:** Overlapped reactions for each class.

| BIGG<br>Reaction ID | Description | PGPR<br>(75 GEMs) | EPP<br>(33 GEMS) | Adjusted<br>p-value |
| --- | --- | --- | --- | --- |
| PGPR |  |  |  |  |
| ACOAD1fr | Acyl-CoA dehydrogenase (butanoyl-CoA) | 54 (72%) | 5 (15%) | 1.4 x 10<sup>-4</sup> |
| DORNabcpp | D-Ornithine transport via ABC system periplasm | 61 (81%) | 6 (18%) | 2.5 x 10<sup>-6</sup> |
| DORNtex | D-ornithine transport via diffusion extracellular to periplasm |  |  |  |
| EX_orn__D_e | Exchange of Ornithine |  |  |  |
| EPP |  |  |  |  |
| CHOLD3 | Choline dehydrogenase | 4 (5%) | 33 (100%) | < 10<sup>-6</sup> |
| FBA2 | D Fructose 1 phosphate D glyceraldehyde 3 phosphate lyase | 2 (3%) | 33 (100%) | < 10<sup>-6</sup> |
| GLUSx | Glutamate synthase NADH2 | 16 (21%) | 33 (100%) | < 10<sup>-6</sup> |
| METRR | Methionine racemase | 14 (19%) | 27 (82%) | 2.5 x 10<sup>-6</sup> |
| MNt2pp | Manganese (Mn<sup>+2</sup>) transport in via proton symport (periplasm) | 7 (9%) | 33 (100%) | < 10<sup>-6</sup> |
| ORNR | Ornithine racemase | 14 (19%) | 27 (82%) | 2.5 x 10<sup>-6</sup> |
| SERR | Serine racemase | 21 (28%) | 28 (85%) | 1.4 x 10<sup>-4</sup> |

Intriguingly, reactions related to D-ornithine are represented in both groups via DORNtex, DORNtex and ORNR. This suggested that D-ornithine is utilized by both the PGPR and the EPP. This result is in line with the pathways annotated in the models. Examination of the utilization pathways shows that D-ornithine is converted to L-proline with 5-Amino-2-oxopentanoate and 1-Pyrroline-2-carboxylate as intermediates through 3 reactions (1) D Amino acid dehydrogenase orn D (DAAD5), (2) 1 Pyrroline 2 carboxylate cyclation (1P2CBXLCYCL) and (3) Delta1 piperideine 2 carboxylate reductase (1P2CBXLR). All three reactions were present in all models. However, the mechanism of D-ornithine acquisition is the key difference between both groups. The PGPRs have transporters annotated with DORNabcpp and DORNtex, thus enabling direct D-ornithine uptake from the medium, while the EPPs were annotated with ornithine racemase (ORNR) that catalyze D-ornithine from L-ornithine instead.

We further examine the annotation quality of two reactions that have their genes annotated, DORNabcpp and ORNR. DORNabcpp involved 4 genes presented in the reference of published *P. putida* KT2440 model (ijN1463) with ‘AND’ logical connective, representing the formation of a protein complex. All 4 genes were identified in our selected models with high similarity of approximately 88% (±6%) identity count. On the other hand, one gene is annotated to ORNR, which is a 3 genes system presented in the reference of published Clostridioides *difficile* 630 model (iCN900) with ’ OR’ logical connective, representing isoenzymes. The annotated gene shows a low similarity of approximately 30% (±1%) identity count. In addition, we also investigate other reactions with close similarity using BLASTP with the custom databases. For the full database, DORNabcpp sequences were identified similarly to Ornithine transport via ABC system (periplasm) and D, L-lysine transport via ABC system periplasm. The annotation results are identical when using the database without the DORNabcpp related genes with the identity score remains at 88% (±7%). In contrast, ORNR sequences were similar to D-serine deaminase and other racemases namely alanine, glutamate, methionine, proline, and serine. The database without the ORNR related genes yield different results. Many proteins were detected with wide range of identity score from 20 to 95%. However, none of them was recognized in any of the models (**Supplementary file S5)**. The results suggested that the D-ornithine transports were annotated with confidence, but ORNR annotation may not be as conclusive.

### Comparative model analyses

GEMs composition reveals that there are metabolic differences between the two classes. To simulate their performance in a biological relevant environment, tomato root exudates corresponding to three stages in the plant growth were selected for simulations ^24^. Additional carbon sources were added to the minimal M9 medium. Day2-medium1: M9 with the addition of oxalate and xylose, Day4-medium2: M9 with the addition of citrate and fructose and Day14-medium3: M9 with the addition of citrate, xylose, and maltose. For each medium and model, we assess their essentiality, which is summarized in Table 2. The table describes the average number of (non-)essential reactions and the total number of (non-)essential reactions of all models along with the variation of both.

**Table 2:** Essentiality analysis using media representing environmental changes.

|  | Average number of reactions |  | Total number of reactions |  |  |
| --- | --- | --- | --- | --- | --- |
|  | Non-essential | Essential | Non-essential | Essential | Variation |
| Day2-medium1 | 2456 ± 193 | 215 ± 7 | 3428 | 159 | 128 |
| Day4-medium2 | 2457 ± 193 | 212 ± 7 | 3431 | 158 | 126 |
| Day14-medium3 | 2457 ± 193 | 214 ± 7 | 3429 | 158 | 128 |

The variation category is particularly interesting as it poses the differences between media which could be translated to the characteristics of each class. For the essential reactions within the variation set of each medium, we obtain the corresponding EC number from the model. The EC numbers were mapped to pathways using KEGG PATHWAY as the reference. This results in a fraction of completeness of each pathway. We performed t-test on the fractions to find significantly different pathways between the two classes (p-value < 0.05) (**Table 3**). As a result, 13 pathways prove enriched in the EPP group where only 2 pathways are enriched in the PGPR group. While most of the pathways seem used in all three media, path:map00564 (Glycerophospholipid metabolism) is missing from Day4-medium2, while path:map01110 (Biosynthesis of secondary metabolites) occurred only in Day14-medium3. Most of the essential pathways are associated with amino acid metabolism which suggests that the EPPs are less flexible in the uptake, utilization, and synthesis of these metabolites.

**Table 3:** Significantly different essential metabolic pathways.

| Map ID | Map description | p-value | Media |
| --- | --- | --- | --- |
| **PGPR** |  |  |  |
| path:map00471 | D-Glutamine and D-glutamate metabolism | 0.025 | 1,2,3 |
| path:map00473 | D-Alanine metabolism | 0.025 | 1,2,3 |
| **EPP** |  |  |  |
| path:map00250 | Alanine, aspartate and glutamate metabolism | 0.010 | 1,2,3 |
| path:map00260 | Glycine, serine and threonine metabolism | 0.025 | 1,2,3 |
| path:map00270 | Cysteine and methionine metabolism | 0.000 | 1,2,3 |
| path:map00340 | Histidine metabolism | 0.004 | 1,2,3 |
| path:map00350 | Tyrosine metabolism | 0.002 | 1,2,3 |
| path:map00360 | Phenylalanine metabolism | 0.004 | 1,2,3 |
| path:map00400 | Phenylalanine, tyrosine and tryptophan biosynthesis | 0.000 | 1,2,3 |
| path:map00401 | Novobiocin biosynthesis | 0.002 | 1,2,3 |
| path:map00564 | Glycerophospholipid metabolism | 0.004 | 1,3 |
| path:map00920 | Sulfur metabolism | 0.025 | 1,2,3 |
| path:map00960 | Tropane, piperidine and pyridine alkaloid biosynthesis | 0.002 | 1,2,3 |
| path:map00997 | Biosynthesis of various secondary metabolites - part 3 | 0.025 | 1,2,3 |
| path:map01110 | Biosynthesis of secondary metabolites | 0.041 | 3 |

We simulated growth flexibility by using flux sampling and differences in the corresponding distributions were evaluated through a Kolmogorov-Smirnov test. In total, 1870 unique reactions were found to carry significantly different fluxes between both groups across all media (**Supplementary file S6**). Reactions were divided into 3 categories: PGPR, EPP and undecided. The undecided group contains reactions in which the reaction direction differs between both classes and no conclusion can be drawn. **Figure 5(a)** shows the frequency of the number of occurrences of the reactions in the comparison. The maximum occurrences are 36 where the reaction is significantly enriched for the entire PGPR group or vice versa. We selected reactions with at least 30 occurrences which result in 107 unique reactions. These reactions are likely to occur in 5 out of 6 of the strains of the same group. The number of reactions were further reduced to 78 by removing reactions appearing as significant in both groups, and in the undecided group. **Figure 5(b)** shows the media occupancy of these 78 reactions and their overlap. Regardless of the media, 57 reactions are shared and could potentially describe the general differences between classes. **Figure 5(c)** shows the 57 overlapping reactions along with their occurrences, media, and significance category. The reactions’ information retrieved from BIGG database are shown in Table **4**.

**Table 4:**
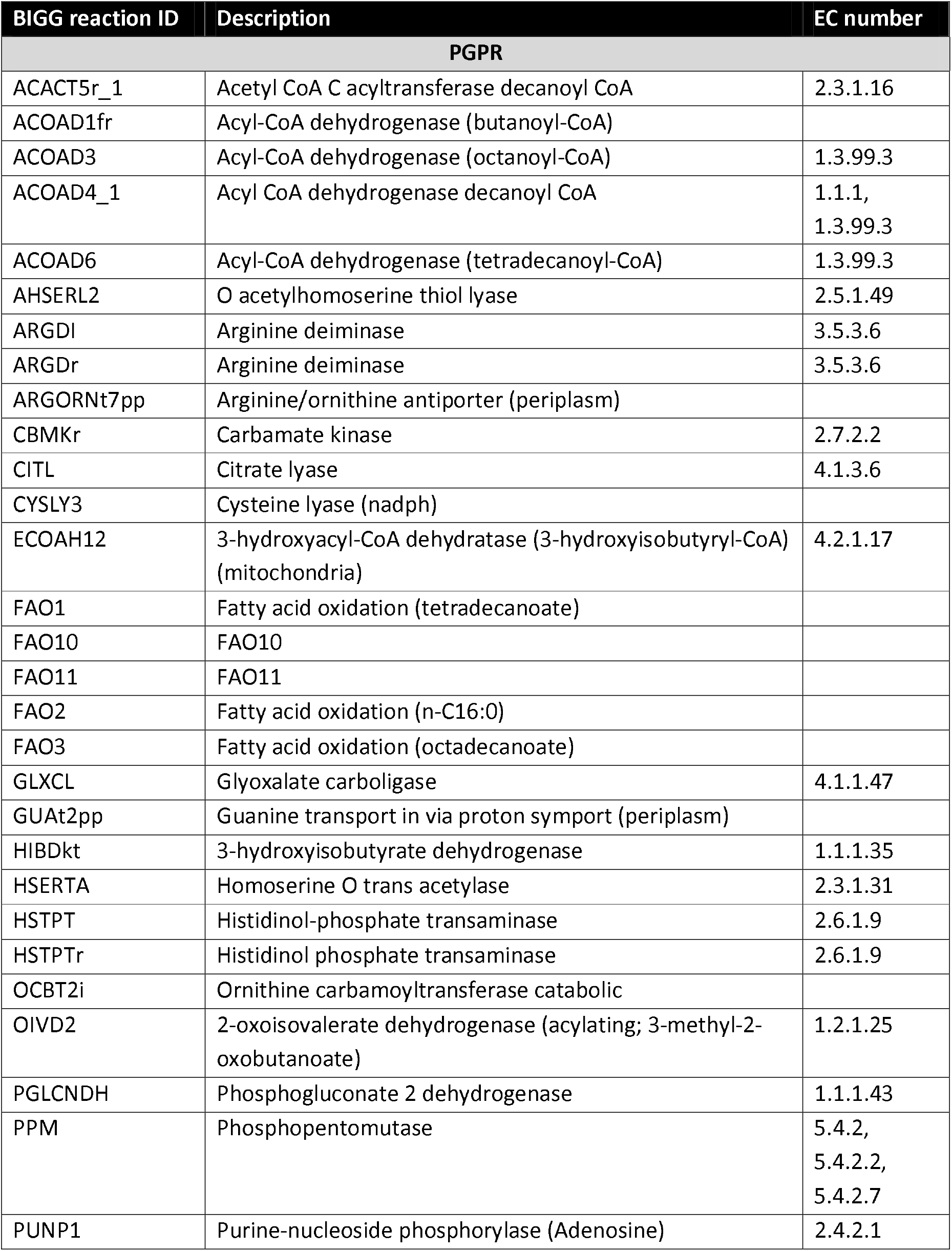
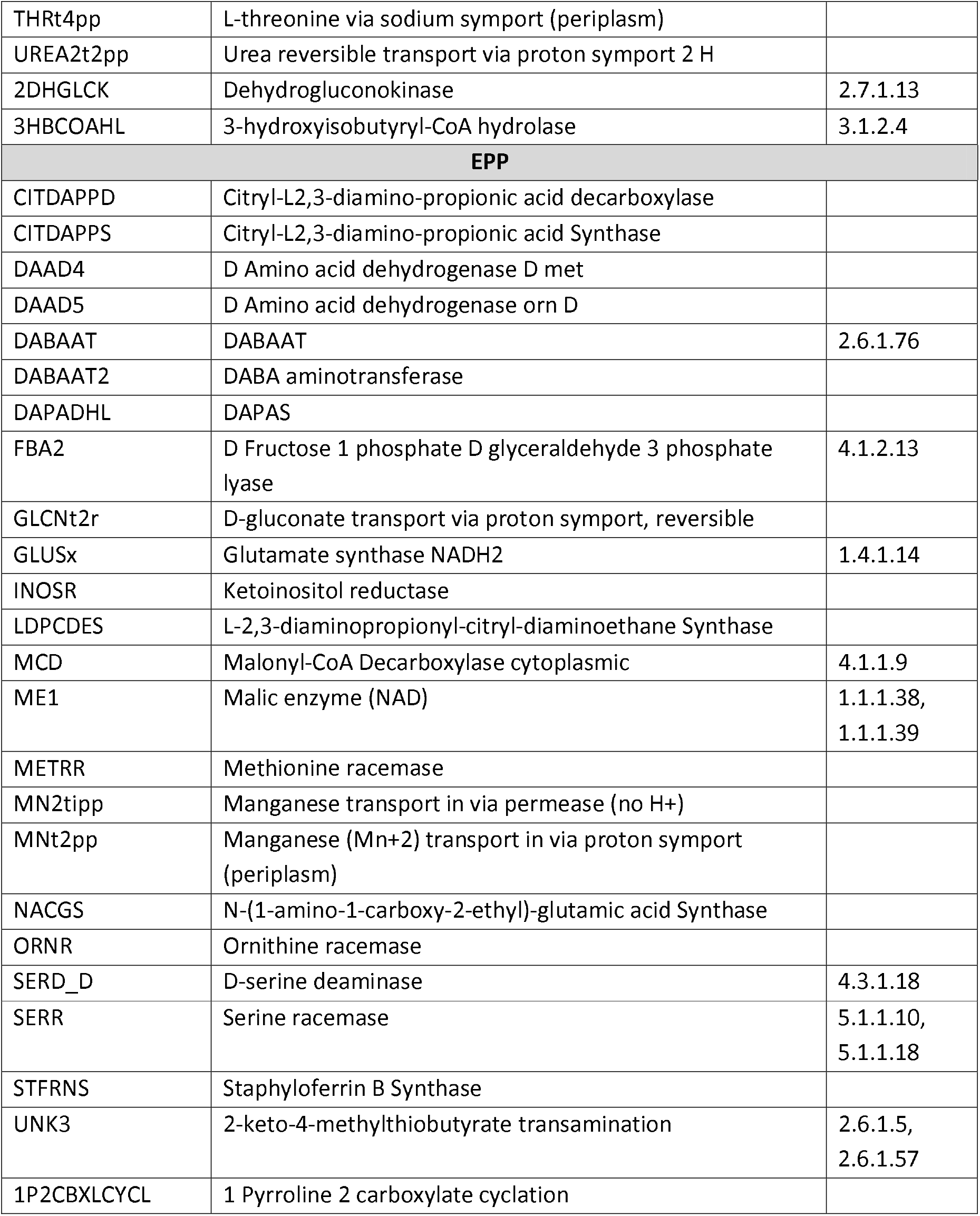
Significant reactions overlapped between all media representing different growth stages and root exudates.

**Figure 5:**
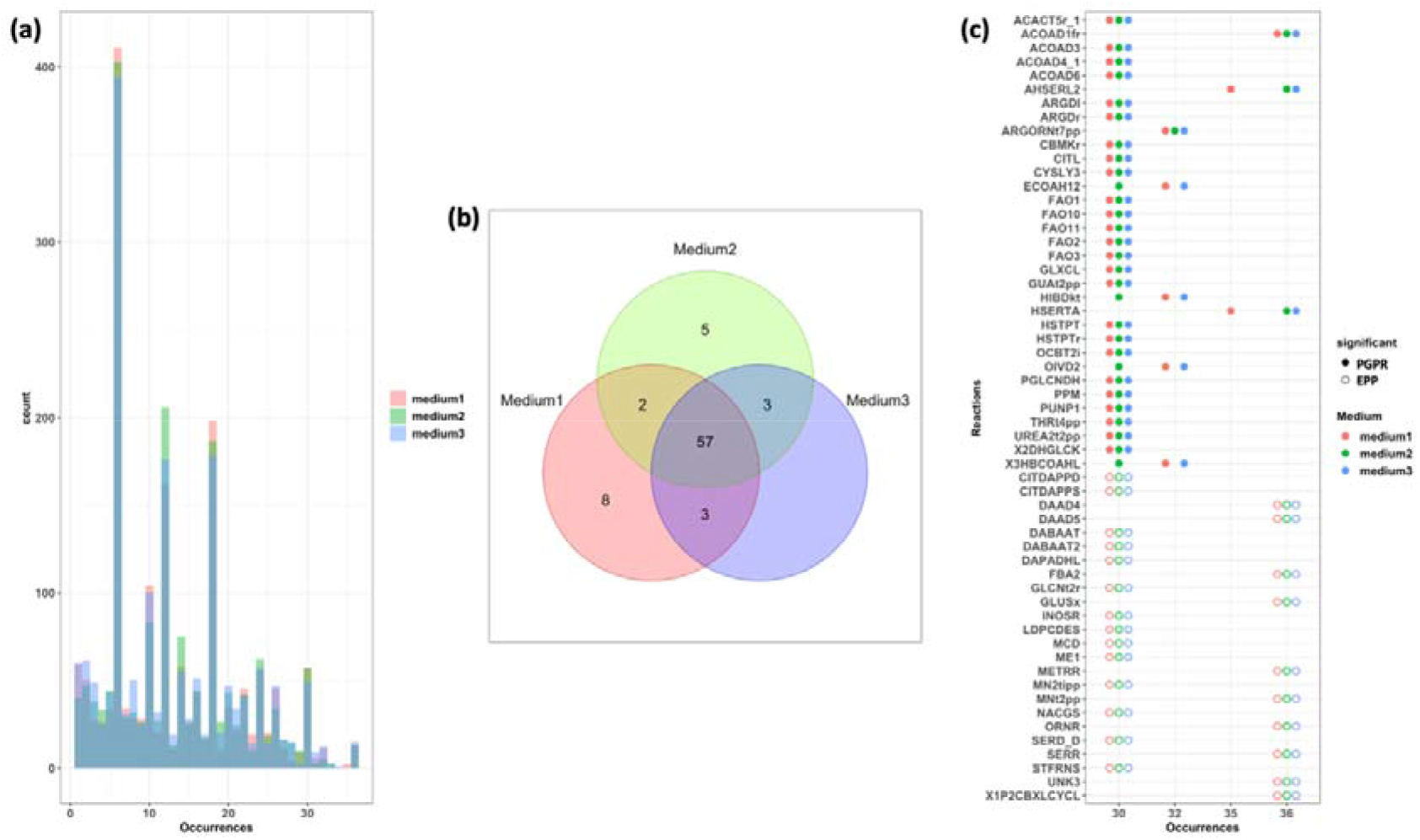
Significantly different reactions between the two classes. Each color represents different media. Red represents Day2-medium1, green represents Day4-medium2 and blue represents Day14-medium3. (a) Histogram shows the number of occurrences of reactions across all media in all the comparisons. (b) Venn diagram of the 78 selected reactions represented in each medium. (c) The 57 overlapped reactions across all media. Significant reactions of each class are indicated by different colored dots. The full dots represent the PGPR group, and the hollow dots represent the EPP group.

A total of 33 and 24 reactions are considered significant overrepresented in the PGPR and the EPP groups, respectively. This information shows different metabolic wiring between the two classes when optimized for growth. The majority of the reactions in the PGPR group are associated with fatty acid oxidation while the EPP mostly consists of racemase reactions and iron acquisition mechanisms.

## Discussion

In this manuscript we demonstrate the usefulness of genome-scale metabolic models to explore the metabolic capacity of organisms in the rhizosphere and gain insights into their potential interactions. The comparative approach on the collection of species belonging to two distinct lifestyle classes, Epiphytic Pathogenic *Pseudomonas* (EPP) and Plant-Growth Promoting Rhizobacteria (PGPR), enables us to identify and characterize unintuitive differences at the metabolic level between the two.

An automatic model construction approach was deliberately chosen because the process to manually curate GEMs is time-consuming and not practical for large-scale comparisons^39–41^. There are several tools for the automation methods and CarveMe was the tool of our choice as it has shown a good performance. Moreover the fact that it is based on a universal model facilitates comparison between models^21,22^. We evaluated the generated model reconstructions and the tool’ s performance by comparing model predictions with actual Biolog phenotype data of a set of strains, which characterize the carbon uptake profile of the organisms. The results show that the performance of the automatic method was acceptable, around 70% even in the absence any manual curation which increases from the original publication of the tool^21^. Additionally, the generated models were composed of proportionally high GPRs with few orphan reactions meaning that the majority of the reactions were supported by evidence of genes. The comparison was performed by considering 55 carbon sources from the 190 measured in our Biolog set as mapping the carbon sources in the model and those in the Biolog data proved rather laborious due to inconsistencies in names^42^. In addition, the comparison disclosed the knowledge gap in the field of automated genome annotation, which results in systematically incorrect predictions, such as acetoacetic acid, glycine, and L-Methionine.

The automated selection of the representative strains also indicated the suitability of our choice of 7 strains to be re-sequenced. Out of the 6 representative PGPRs, three were among the re-sequenced strains, suggesting that our selection covers a broad phylogeny range and mode of actions for the PGPRs *Pseudomonas*. Outliers from figure 4, *P. stutzeri* A1501 123 and *P. cichorii* JBC1 2922, were also included to maximize the range of the represented groups, PGPRs and EPPs respectively.

Both the construction and the simulation of GEMs highlighted the metabolic differences between the two plant-related phenotypes, the PGPRs and the EPPs. Differences were observed in their potential to transport compounds such as amino acids, sugars, or metal ions, in and out of the cellular environment (transport reactions), these signify the compositional differences between both classes and can highlight their distinct behavior. The PGPRs were enriched with various amino acid related transporters whereas reactions related to amino acid synthesis were often found to be essential in the EPPs. This suggests that PGPRs *Pseudomonas* are able to import amino acids from their environment whereas the EPPs can only rely on intracellular synthesis and have limited uptake capabilities. This suggests a critical role of amino acids emitted by the host plant to affect community composition, as has been previously studied in E. coli^43,44^. On the other hand, the abundance of sugar transporters in the EPPs points to their nutrient preferences, such as galactose, xylose, and sucrose. It appears associated to their parasitic lifestyle, which is more dependent than PGRPs on carbon from the host for proliferation, as it has been shown in *Xanthomonas oryzae* and *Pseudomonas syringae*^45^. This information on preferred carbon sources can be used to develop rhizosphere management strategies aiming to exclude pathogens or to improve numbers of beneficial organisms^46–48^.

Differences in D-ornithine acquisition mechanisms were observed when comparing both groups. While this metabolite is relevant for the metabolism of organisms in both groups, differences in the acquisition mechanisms were identified. The PGPR organisms can potentially take up D-ornithine from the environment whereas the EPP appear to be capable of intracellular conversion from L-ornithine using a racemase reaction. With L-proline being the sole final product, the evidence implies that the PGPR class would benefit from an environment with limited L-proline and L-ornithine while supplied with D-ornithine. Additional analysis suggests other D-amino acid and racemases could share the same characteristics related to differential uptake and utilization mechanisms, for example D-lysine and D-arginine^49,50^. D-amino acids have been found abundantly in soil inhabited by microbiomes with annotated racemases^51^. These substrates can be taken up by both the microorganisms and the hosts. However, it appears to be more relevant for microbial growth than plant growth as D-amino acids were found prominently in bacterial cell walls while they inhibit growth of some plants^52–55^. This may suggest another beneficial effects of the microbiome, which would be consumption of D-amino acids and their removal from the environment.

In addition to the differences in the abovementioned transports, the two bacterial classes have different internal metabolic wiring shown through the flux sampling analysis when optimized towards the optimal growth in different conditions. The flux sampling comparison reveals that, in general, the PGPRs have more active fatty acid metabolisms whereas the EPPs activate pathways related to the racemization and iron and metal acquisition mechanisms. In plants, fatty acids are markers of both biotic and abiotic stresses^56,57^. Since these fatty acids are transferable between the plant and the rhizobacteria^58^, we hypothesize that the activation of these pathways in PGPRs to be a potential signal for a reinforcement from the host in combination with the amino acid secretion. On the other hand, racemization and iron scavenging are prominent in the EPPs and both functions deplete fundamental substrates from the environment, L-amino acids, and metals respectively, which are essential and competitive substrates for both the plant and the microorganisms^59,60^. We emphasise that iron is known to be essential to all living organisms, which also represented by 23 common iron-related reactions that occur in all models (**Supplementary file S7**), however EPPs seem to have more efficient mechanisms to overcome this limitation. In addition, the iron-limited environment negatively affects the production of the crops and could possibly alter the behavior of the beneficial microbes, like *Pseudomonas fluorescens* BBc6R8^61,62^. Similar conclusions can be drawn using Genome Properties (GPs) and the dynamic nature of the simulation in GEMs further support the results^25^.

## Conclusion

It has been shown that genome-scale constraint-based metabolic modelling is a viable approach to represent the metabolic capacity of an organism. Here, we want to emphasize that GEMs can also be used to compare metabolic spaces and gain insights into differences in metabolic behavior and implications for the environments where these microbes thrive. In addition, the validated automation method enables comparative analysis and potentially broadens the scope of the study into modeling the entire microbial community. The model allowed us to explore an organism using both composition and simulation methods, which are the composition and the simulation of the model respectively. Both methods were able to differentiate between PGPR and EPP *Pseudomonas* strains. Some differences could be used to explain the underlying mechanisms of the distinct lifestyle between two classes, such as the fatty acid and iron acquisition mechanisms, while other differences, such as amino acid and sugar transports, could be incorporate into the development of the rhizosphere management strategies to precisely assist beneficial microbiome while reducing the pathogen activities.

## Acknowledgements

WP is financially supported by a Royal Thai Government Scholarship, Thailand. TL acknowledges the support by the Dutch Ministry of Economic Affairs in the Topsector Program “Horticulture and Starting Materials” under the theme “Plant Health” (project number: TU 16022) and its partners (NAK, Naktuinbouw and BKD). PS and MSD acknowledge the Dutch national funding agency NWO, and Wageningen University and Research for their financial contribution to the Unlock initiative (NWO: 184.035.007).

## Author Contributions

All authors participated in the conception and design of the study. A.D.v.D. and T.A.J.L. provided the phenotype microarray (Biolog) data and phenotypic classification of the strains. W.P. performed the computational analyses. W.P. wrote the original draft of the manuscript. W.P., A.D.D., T.A.J.L., VMdS, PJS and M.S.-D., contributed to the writing, review, and editing of the manuscript.

## Data availability

The datasets and computer code produced in this study are available in Gitlab at https://gitlab.com/wurssb/pseudomonas-gems

## Notes

https://gitlab.com/wurssb/pseudomonas-gems

### Competing Interest Statement

The authors have declared no competing interest.

