## Supplementary material for "Comparative genome-scale constraint-based metabolic modeling reveals key lifestyle features of plant-associated *Pseudomonas* spp": Supplementary file S1 - script example.py

```
# !/usr/bin/env python
# -*- coding: utf-8 -*-

"""
Author: Wasin Poncheewin
		pFBA implementation from Rik van Rosmalen, util.py
		https://gitlab.com/wurssb/Modelling/sampling-tools/-/blob/master/sampling_tools/util.py
"""

import cobra
import cobra.test
from cobra import Model, Reaction, Metabolite
import os
from os import listdir, makedirs
from os.path import isfile, join
import sys
import pandas as pd
import itertools
from cobra.flux_analysis import (
    single_gene_deletion, single_reaction_deletion, double_gene_deletion,
    double_reaction_deletion, flux_variability_analysis, production_envelope)
from cobra.sampling import sample
from cobra.util.solver import linear_reaction_coefficients

def add_pFBA_constraint(model, objective, objective_fraction=1, flux_fraction=1):
    """Add a pFBA flux minimality constraint to the model. Updates the model in-place.

    :param model: The cobra model
    :type model: cobra.Model
    :param objective: The objective that should be maximized,
    in order to determine the objective bound for the flux sum minimization.
    :type objective: cobra.Reaction
    :param objective_fraction: pFBA minimum objective fraction, should be 1<= and defaults to 1.
    :type objective_fraction: float, optional
    :param flux_fraction: pFBA maximum flux fraction, should be >=1 and defaults to 1.
    :type flux_fraction: float, optional
    :return: The model with the flux minimality constraint added.
    :rtype: cobra.Model
    """

    def recursive_sum(x):
        """Recursive version of sum.

        Sum operations are slow on variables. By doing it recursively, we can use
        2log(length(x)) additions instead of length(x).
        We could exceed the recursion depth, but it is not very likely with normal model sizes.
        """
        length = len(x)
        if length == 1:
            return x[0]
        elif length == 2:
            return x[0] + x[1]
        else:
            split = length // 2
            return recursive_sum(x[split:]) + recursive_sum(x[:split])

    # Calculate flux sum for the objective.
    flux_sum_value = cobra.flux_analysis.pfba(model, objective_fraction, objective)

    # Create a total flux variable (fw + rev fluxes, same as pFBA does it in Cobra.)
    variables = ((r.forward_variable, r.reverse_variable) for r in model.reactions)
    sum_variable = recursive_sum(list(itertools.chain.from_iterable(variables)))
    # Create and add the constraint to the model.
    constraint = model.problem.Constraint(
        expression=sum_variable,
        name="total_flux_sum_bound",
        lb=0,
        ub=flux_sum_value.objective_value * flux_fraction,
    )
    model.add_cons_vars([constraint])
    return model

if __name__ == '__main__':

	# prepare output folder
	if not os.path.exists('output_folder/'):
		os.makedirs("output_folder", exist_ok=True)

	# essentiality analysis folder
	if not os.path.exists('output_folder/srd/'):
		os.makedirs("output_folder/srd", exist_ok=True)

	# flux sampling analysis folder
	if not os.path.exists('output_folder/flx_pfba/'):
		os.makedirs("output_folder/flx_pfba", exist_ok=True)

	outFile = open('growth.tsv', 'w')
	outFile.write('Organisms\tm9_growth\tadjusted_medium\n')
	
	folder = '../'

	# list all models
	onlyfiles = [f for f in listdir(folder) if isfile(join(folder, f))]
	
	for f in onlyfiles:
		if f.endswith('.xml'):
			print (f)
			modelname = folder + f
			model = cobra.io.read_sbml_model(modelname)

			m9_growth = model.optimize()
			print ("Growth:" + str(m9_growth.objective_value))
			
			print (model.medium)

			# additional substrates for medium1
			model.reactions.get_by_id('EX_oxa_e').lower_bound = -15
			model.reactions.get_by_id('EX_xyl__D_e').lower_bound = -11

			# additional substrates for medium2
			model.reactions.get_by_id('EX_cit_e').lower_bound = -5
			model.reactions.get_by_id('EX_fru_e').lower_bound = -9.17

			# additional substrates for medium3
			model.reactions.get_by_id('EX_cit_e').lower_bound = -5
			model.reactions.get_by_id('EX_xyl__D_e').lower_bound = -5
			model.reactions.get_by_id('EX_malt_e').lower_bound = -2.5

			add_growth = model.optimize()
			print ("Growth:" + str(add_growth.objective_value))

			print (model.medium)

			outFile.write(f.replace('.xml','') + '\t' + str(m9_growth.objective_value) + '\t' + str(add_growth.objective_value) + '\n')
			
			# essentiality analysis 
			single_reaction_deletion(model).to_csv('output_folder/srd/' + f.replace('.xml','.csv'))

			model = add_pFBA_constraint(model, model.objective)

			# flux sampling analysis
			flx = sample(model, 10000)
			flx.to_csv('output_folder/flx_pfba/' + f.replace('.xml','_flx.csv'))

	outFile.close()
```
