## Supplementary material for "Comparative genome-scale constraint-based metabolic modeling reveals key lifestyle features of plant-associated *Pseudomonas* spp": Supplementary file S4 - Venn diagram.docx

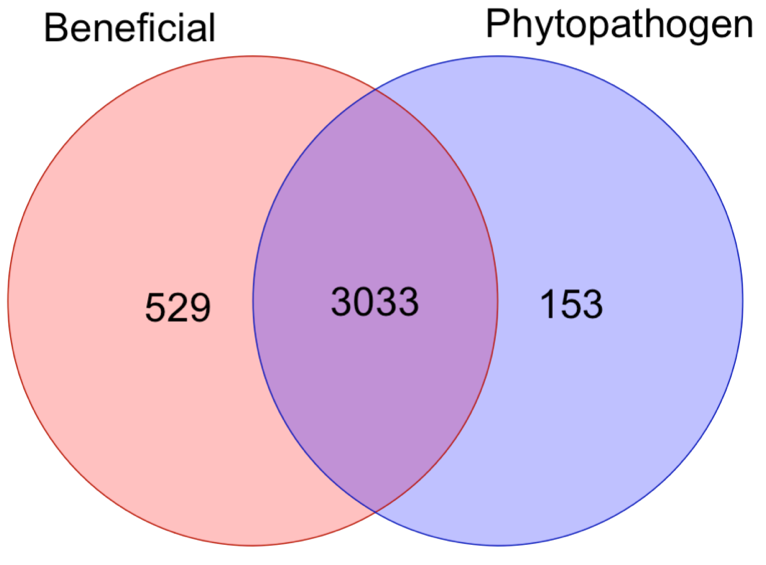

**(a)**

**(b)**

**(c)**

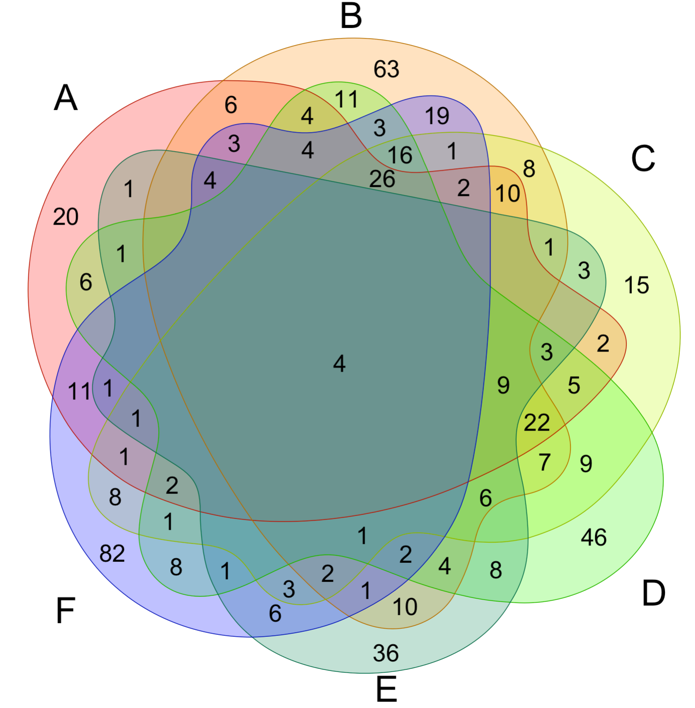

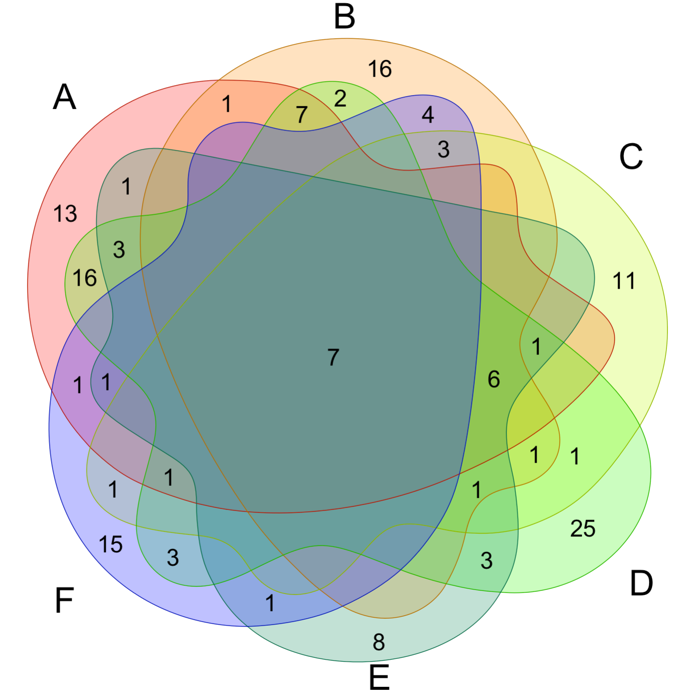

**7**

**4**

**Supplementary figure S4: The Venn diagram representing overlapping and unique reactions compared between groups of 12 representative strains.** (a) Comparison between the selected group of beneficials and the pathogens. (b) Comparison within the 6 selected beneficial strains. The candidates starting clockwise from the A symbol are *P. sp*. UW4, *P. chlororaphis* Phz24, *P. fluorescens* WCS374, *P. jessenii* RU47, *P. rhizosphaerae* DSM 16299, *P. stutzeri* A1501. (c) Comparison within the 6 selected pathogenic strains. The candidates starting clockwise from the A symbol are *P. syringae* pv lapsa ATCC 10859, *P. viridiflava* CFBP 1590 isolate E12-5, *P. syringae* B13-200, *P. savastanoi* pv phaseolicola 1448A, *P. cerasi* isolate Sour cherry Prunus cerasus symptoma, *P. cichorii* JBC1.
