## Supplementary material for "Comparative genome-scale constraint-based metabolic modeling reveals key lifestyle features of plant-associated *Pseudomonas* spp": Supplementary file S5 - D-ornithine annotation.docx

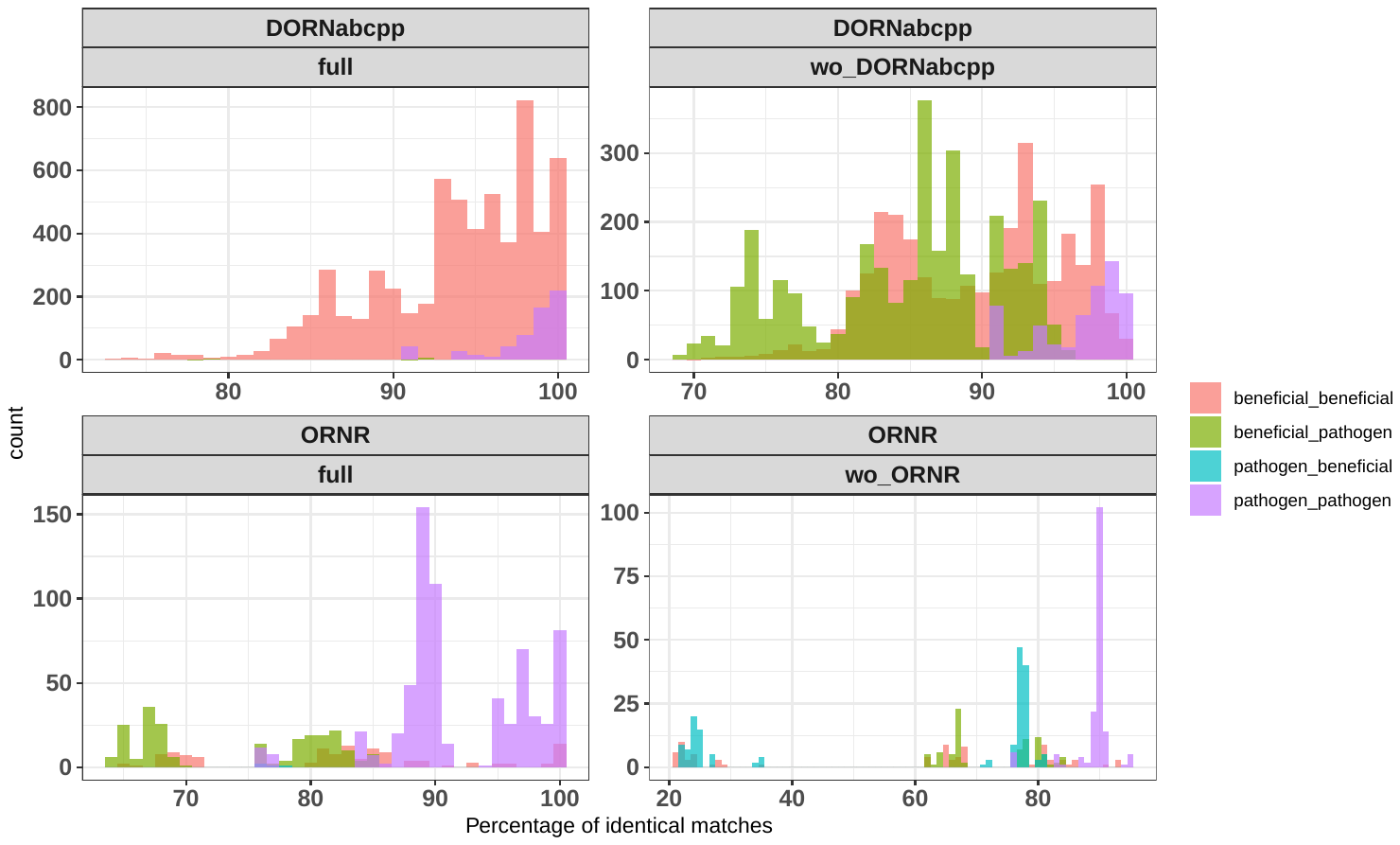

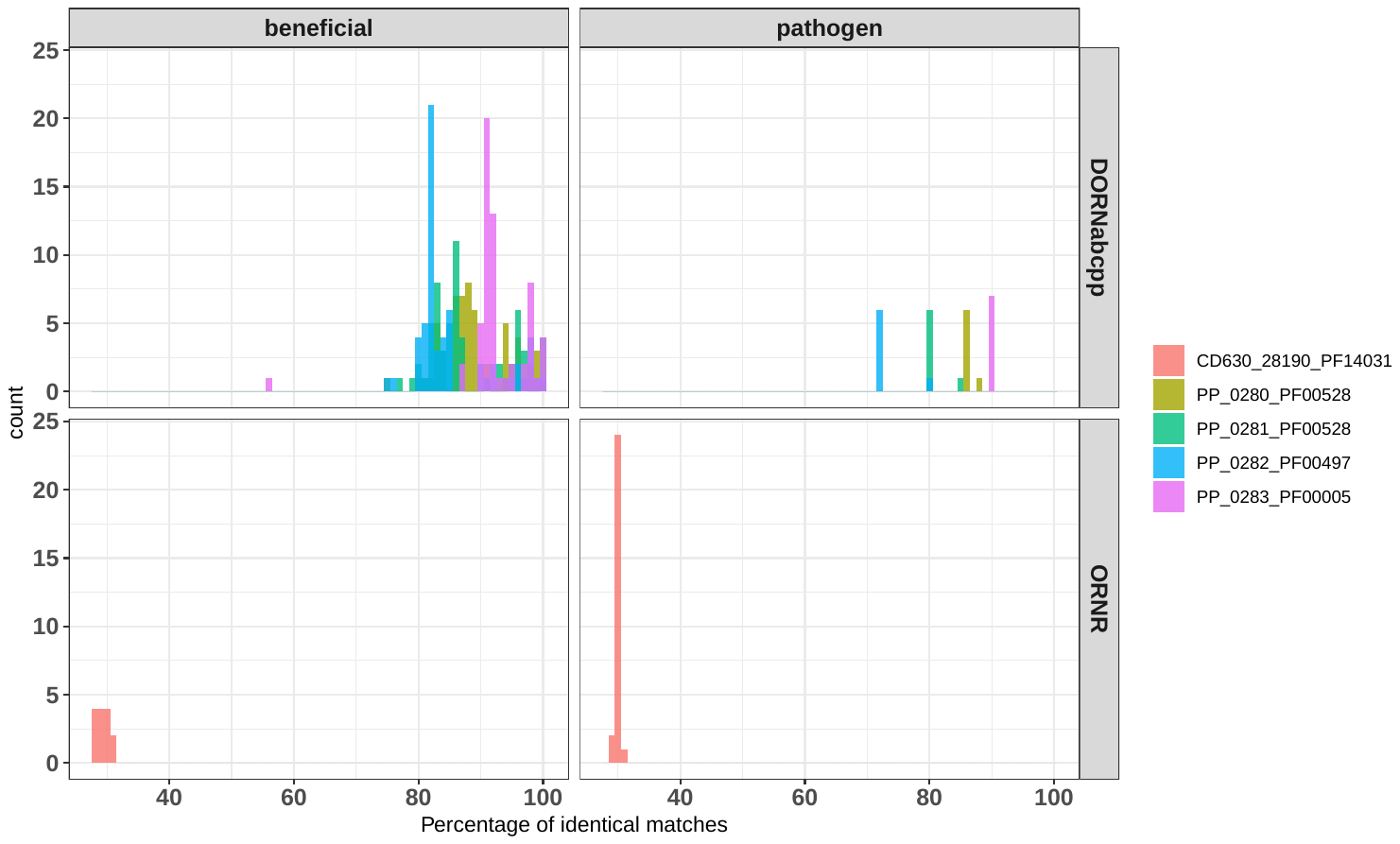

**(b)**

**(a)**

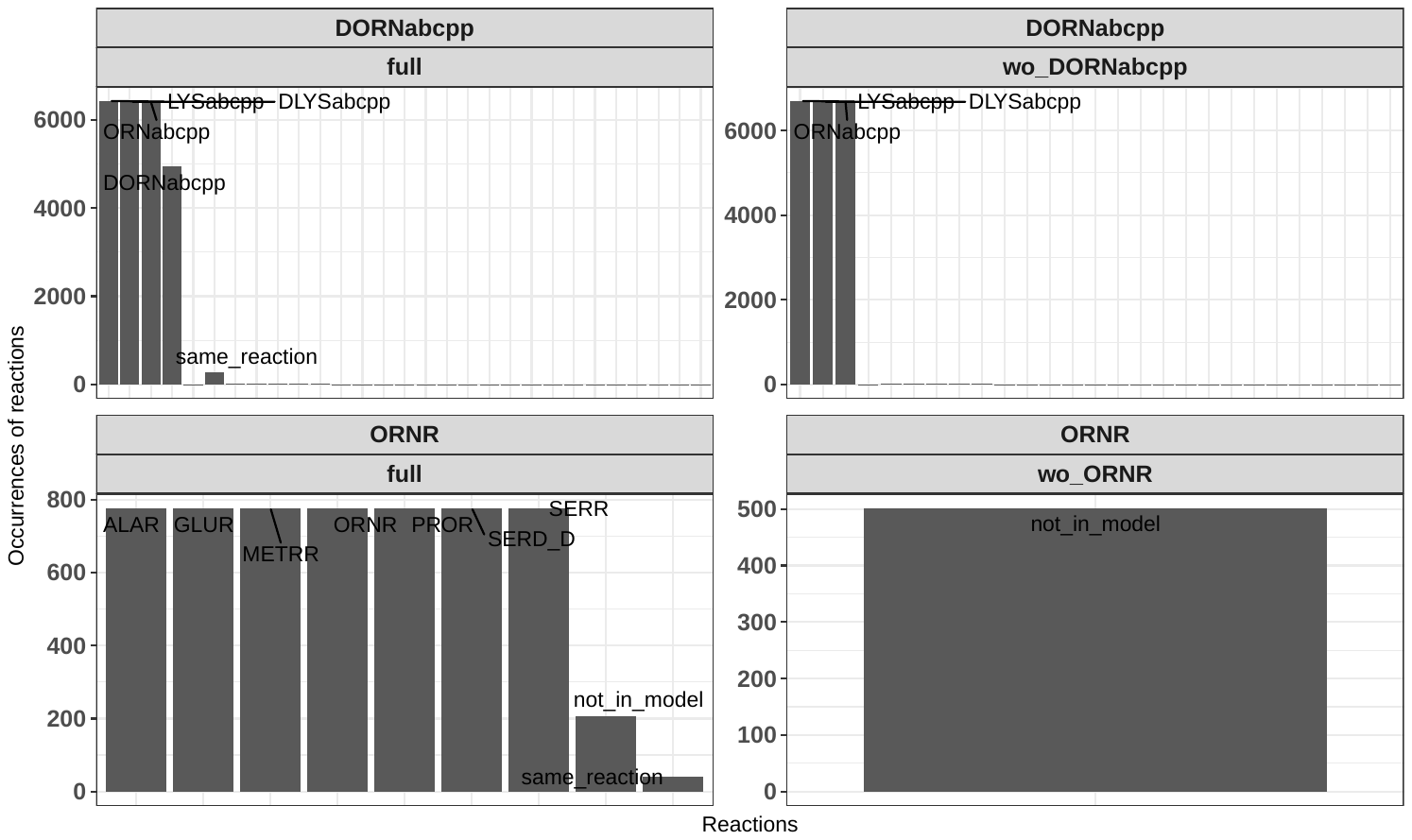

**(c)**

**Supplementary figure S5: The annotation of D-ornithine related reactions: DORNabcpp and ORNR.** (a) The quality of the annotation within the models of the reaction related genes shown by percentage of identical matches. The red color represents the gene for ORNR, and the other colors represents the 4 genes related to DORNabcpp. (b) The percentage of identical matches from BLASTP results of the corresponding genes against the custom databases. “full” represents the full set of genes including the gene in consideration itself, where “wo_” is the database without the gene in consideration. We separated the gene sequences found in beneficials and pathogens, which represents by different colors. (c) The description of the matches.
